# Robust and highly efficient transformation method for a minimal Mycoplasma cell

**DOI:** 10.1101/2024.09.23.614646

**Authors:** Masaki Mizutani, John I Glass, Takema Fukatsu, Yo Suzuki, Shigeyuki Kakizawa

## Abstract

Mycoplasmas have been widely investigated for their pathogenicity, as well as for genomics and synthetic biology. Conventionally, transformation of mycoplasmas was not highly efficient, and due to the low transformation efficiency, large amounts of DNA and recipient cells were required for that purpose. Here we report a robust and highly efficient transformation method for the minimal cell JCVI-syn3B, which was created through streamlining of the genome of *Mycoplasma mycoides*. When the growth states of JCVI-syn3B were examined in detail by focusing on such factors as pH, color, absorbance, CFU, and transformation efficiency, we found that the growth phase after the lag phase can be divided into three distinct phases, of which the highest transformation efficiency was observed during the early-exponential growth phase. Notably, we attained the transformation efficiency up to 4.4 × 10^-2^ transformants per cell per μg plasmid DNA. We developed a method to obtain several hundred to several thousand transformants with less than 0.2 mL of cultured cells and 10 ng of plasmid DNA. Moreover, we established a transformation method using frozen stock of transformation-ready cells. These procedures and information could simplify and enhance the transformation process of minimal cells, facilitating advanced genetic engineering and biological research using minimal cells.

**IMPORTANCE:** Mycoplasmas are parasitic and pathogenic bacteria for many animals. They are also useful bacteria to understand cellular process of life and for bioengineering because of their simple metabolism, small genomes and cultivability. Genetic manipulation is crucial for these purposes, but transformation efficiency in mycoplasmas is typically quite low. Here we report a highly efficient transformation method for the minimal genome mycoplasma JCVI-syn3B. Using this method, transformants can be obtainable only 10 nanograms of plasmid DNA, which is around one-thousandth amount required for traditional mycoplasma transformations. Moreover, we established a convenient method using frozen stocks of transformation-ready cells. These improved methods play a crucial role in further studies using minimal cells.

## INTRODUCTION

The bacteria belonging to the class Mollicutes, which represent *Mycoplasma*, *Spiroplasma*, *Ureaplasma*, *Candidatus* Phytoplasma and others, are parasitic and occasionally commensal bacteria that are characterized by small genome and lacking peptidoglycan cell wall (1, 2). Many bacteria in the class Mollicutes are pathogenic species of various host organisms and causes not only health problem but also large economic loss (3–5). Mollicutes bacteria are also important for bacterial genome research due to their small genomes and cultivability. Numerous milestone studies in bacteria have been conducted using mycoplasmas. For example, scientific accomplishments in mycoplasmas include: the second bacterium whose entire genome was sequenced (6, 7), the first bacterium with a successful genome transplantation (8), whose genome was fully synthesized and transplanted (9), and minimized (10). Additionally, several projects using *Mycoplasma* species as bacterial chasses are currently progressing (11–13).

JCVI-syn3.0 (a minimal cell) is a bacterium with the smallest genome in culturable bacteria, whose 531-kbp genome encodes only 473 genes (10). JCVI-syn3.0 was created based on *Mycoplasma mycoides*; it grows well and is capable of transformation. Various tools have been developed for JCVI-syn3.0, such as genome insertion tools using Cre-*lox* recombination system (14–16), gene knockdown systems using inducible promoters and CRISPR-interference (16), and *oriC* plasmid systems (14, 16). JCVI-syn3A and JCVI-syn3B were derivative strains with 19 and 20 additional genes compared to JCVI-syn3.0, respectively (17, 18). These strains showed relatively normal cell division and morphology, and they grow fast, making them useful for research. JCVI-syn3B is nearly identical to strain JCVI-syn3A, differing only in the presence of a dual *loxP* site landing pad in its genome for DNA introduction. Various studies using these minimal cells were reported so far, such as reconstruction of spiral cell shape and swimming motility of *Spiroplasma* (19), reconstruction of attachment ability and pathogenicity of *Ureaplasma* to human cells (20), understanding of cellular metabolic networks through simulations (21, 22), analysis of cell division mechanisms (17), laboratory evolution experiments (18, 23), and interactions with human cells (13, 20).

Transformation and gene introduction are crucial for many purposes, including the expression and purification of proteins, plasmid construction, production of biological compounds, and elucidation of gene function and cellular biology. In many mycoplasmas, lots of transformation systems and methods have been reported so far. However, the transformation of mycoplasmas is generally inefficient and for some species still not achieved. In most cases, a large amount of plasmid DNA are required, around 10 micrograms, and a large volume of freshly cultured cells is also needed. Despite these measures, transformation efficiency remains low (≤ 10^-7^ transformants/cells/µg plasmid DNA) (24–27). This low efficiency can be a significant impediment to mycoplasma genetic engineering. To establish mycoplasmas as one of the model organisms, to contribute to the elucidation of fundamental biology using mycoplasmas, and to utilize mycoplasmas for various applications as a chassis in synthetic biology, it is essential to develop advanced genetic engineering tools along with effective gene introduction methods (28–30), thereby improving utility of mycoplasmas. Additionally, detailed descriptions of basic biological information, such as growth rate, growth stage, number of living cells in culture, transformation efficiency, would also be necessary.

Here, we report a robust and highly efficient transformation method for JCVI-syn3B. We successfully developed a method to obtain several hundred transformants using less than 0.2 mL of bacterial culture and 10 ng of plasmid DNA. We also developed a method for frozen cell stock in a transformation-ready state, enabling their use similar to competent cells of *Escherichia coli*. During this process, we discovered that the exponential phase of JCVI-syn3B can be divided into two distinct stages, which significantly affects transformation efficiency. Additionally, we report basic data on the pH, color of medium, light absorbance, and number of viable cells in culture (colony forming unit: CFU). These methods and data obtained in this study would contribute to the further use of the minimal cells.

## MATERIALS AND METHODS

### Bacterial strains and culture conditions

JCVI-syn3B (GenBank: CP146056.1) was statically grown in SP4 medium at 37°C (31). The recipe of SP4 medium is shown in Supplementary Table S1. *E. coli* strain DH5α (New England Biolabs, Massachusetts, U.S.) was grown in LB medium (5 g/L Yeast extract, 10 g/L Tryptone, 5 g/L NaCl) containing 25 μg/mL Zeocin at 37°C with shaking.

### pH measurement and spectrometry

Various coloring SP4 media were prepared by serial addition of HCl. pH of each SP4 medium was measured by a pH meter (LAQUAtwin pH-22B; HORIBA, Kyoto, Japan). Color of medium was pictured by a scanner (GT-X830; Epson, Tokyo, Japan). Light absorbance at 560-nm wavelength was measured by a plate reader (Absorbance96; byonoy, Hamburg, Germany) and normalized the absorbance value of intact SP4 medium as 0. Plot illustration and linear fitting were performed using IGOR Pro 9 (WaveMetrics, Oregon, U.S.). For cell growth measurement, 10 μL frozen stock of JCVI-syn3B cells (5.0 × 10^5^ CFU) was inoculated into 190 μL of SP4 medium in 96-well plate. Changes of light absorbance at 560 nm were monitored by the plate reader for 2 days with 30-min intervals at 37°C.

### CFU measurement

Cell cultures at various growth stages were 10^3^‒10^4^-fold diluted, plated onto SP4 agar plate and incubated at 37°C for four to five days. Number of colonies were counted through a stereo microscope (M125C; LEICA, Wetzlar, Germany) and calculated colony forming unit per mL of culture.

### Plasmid construction and purification

To measure transformation efficiency, we constructed simple plasmids for the minimal cell JCVI-syn3B. Both the Cre-*lox* system (Landing pad system) and *oriC* plasmid system have been reported in minimal cells (16), so we decided to measure the transformation efficiency using both systems. Starting with pSD073 (11.8 kb, *oriC* plasmid for CRISPRi) and pSD079 (10.9 kb, landing pad plasmid for CRISPRi) (16), we removed unnecessary regions except for the puromycin resistance gene. PCRs were performed with two primers, Del_CRISPRi-mCh_F1 (5’-AAT TCA AAA AAT AAG GAC TGA GCT AGC TGT CAA AGA TC-3’) and Del_CRISPRi-mCh_R1 (5’-CTA GCT CAG TCC TTA TTT TTT GAA TTA AGT ATT AAA TAA GTG-3’), and the products were recircularized by self-closure, generating plasmids pSD127 (*oriC* plasmid) and pSD128 (landing pad plasmid). Sanger sequencing revealed an unintended nonsense mutation within the puromycin resistance gene of pSD127. This nonsense mutation would inactivate the puromycin resistance gene, so we fixed this mutation by amplifying the plasmid with the Puro_Stop_Corr_F1 (5’-AA AAA TAA TTC TTG TAA TTC AGT AAC TCT TTC AAT ATG TC-3’) and Puro_Stop_Corr_R1 (5’-CTG AAT TAC AAG AAT TAT TTT TAA CTA GAG TTG GTT TAG-3’) primers and recircularizing the products, generating plasmid pSD131 (*oriC* plasmid). The sequences of both plasmids were confirmed by Sanger and nanopore methods. The maps of both plasmids are shown in Fig. 1. pSD128 is a Cre-*lox* recombination plasmid (Landing pad plasmid). Due to the action of Cre recombinase encoded on the plasmid, DNA recombination occurs between the *loxP* sites on the plasmid and on the JCVI-syn3B genome (so called landing pad), introducing the puromycin resistance gene into the genome. pSD128 does not replicate in mycoplasma cells. pSD131 is an *oriC*-plasmid with the replication origin of the genome, and it could replicate within the minimal cell.

**FIGURE 1.**
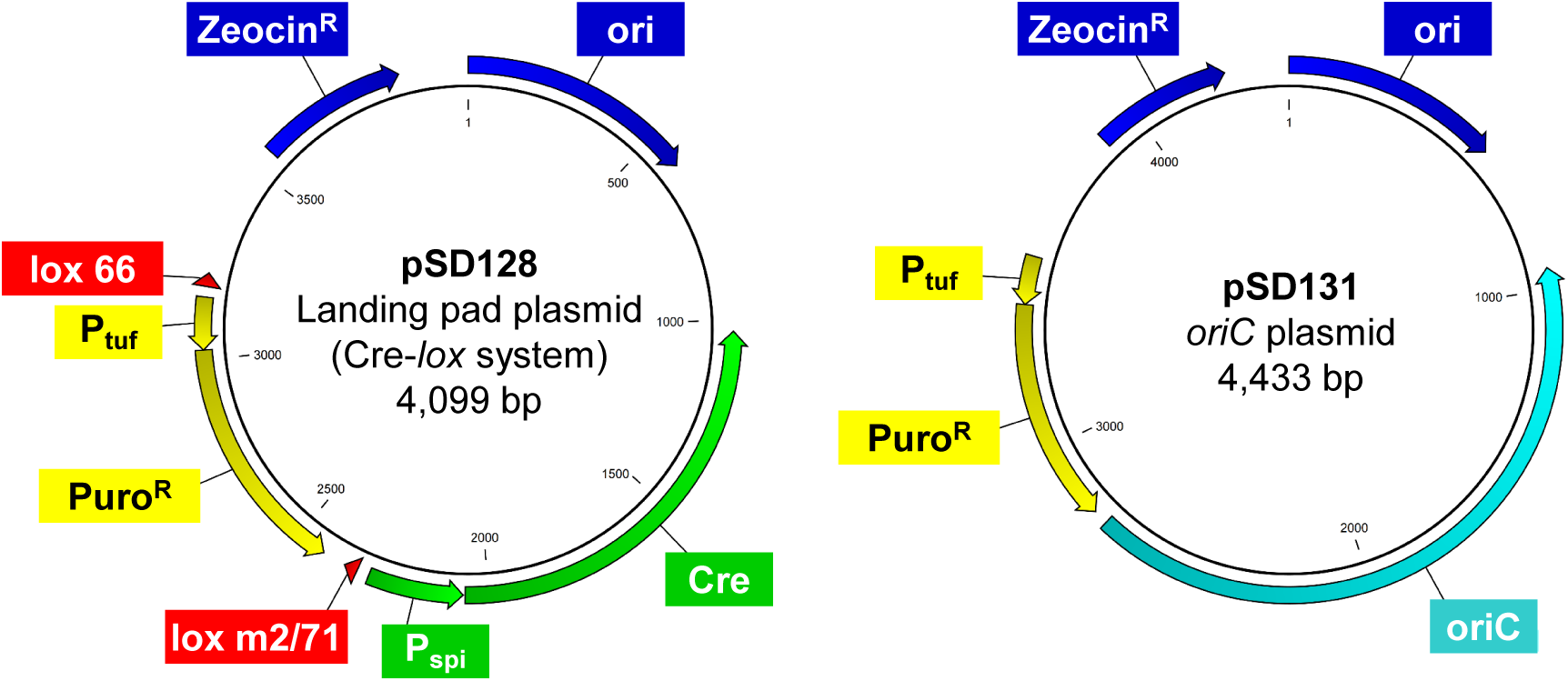
Maps of Plasmids used in this study. pSD128 and pSD131 are a Cre-*lox* recombination plasmid and an *oriC* plasmid, respectively. pSD128 could not replicate in *Mycoplasma* but could introduce the puromycin resistance gene into the genome of JCVI-syn3B. pSD131 could replicate in the minimal cell. Detailed information is described in Materials and Method. ZeocinR: the zeocin resistant gene that is functional in *E. coli*. Mm_oriC: replication origin of chromosome (genome) of *Mycoplasma mycoides* subsp. *capli*, including *dnaA* gene. Ori: replication origin derived from pUC19 plasmid vector in *E. coli*. PuroR: the puromycin resistant gene that is functional in *Mycoplasma*. lox 66 and lox m2/71: *loxP* sites. Ptuf and Pspi are promoter sequences from mycoplasma EF-Tu and Spiroplasma spiralin genes, respectively.

*E. coli* strain DH5α harboring pSD128 and pSD131 plasmids were grown to an optical density at 600 nm = 3 and 5, respectively. Plasmids were isolated from 1 mL of individual cultures using a plasmid purification kit (QIAprep Spin Miniprep Kit; Qiagen, Hilden, Germany) according to manufacturer’s instructions. Concentrations of purified plasmids were measured by a fluorometer (Qubit 4 Fluorometer; Invitrogen, Massachusetts, U.S.).

### PEG-mediated transformation

Polyethylene glycol (PEG)-mediated transformation method of synthetic mycoplasmas was modified based on the previously described methods (19, 26). Summary of transformation procedures developed in this study is shown in Supplementary Figure S1. One mL of culture was centrifuged at 9,000 g for 8 min at 22°C. The pellet was suspended with 1 mL of S/T buffer (0.5 M sucrose, 10 mM Tris-HCl; pH 6.5), centrifuged again, suspended with 140 μL of 0.1 M CaCl_2_ solution and incubated for 30 min on ice. Plasmid DNA solution was mixed with 17 μL of cell suspension in flat bottom 2-mL tube and incubated for 15 min at room temperature (RT). Then, 133 μL of 70% PEG6000 (Averaged molecular weight = 5000‒7000, Sigma-Aldrich, Missouri, U.S.) dissolved in S/T buffer was added, mixed by gentle pipetting using wide bore 200 μL tip (INA OPTIKA, Osaka, Japan). After 2 min incubation at RT, 800 μL of S/T buffer was added and mixed by inverting the tube until completely mix PEG and S/T buffer. The suspension was centrifuged at 10,000 g for 15 min at 10°C. The pellet was suspended with 500 μL of SP4 medium without antibiotics by inverting the tube and incubated at 37°C for 1‒3 hours. Then, the suspension was 10^0^‒10^2^-fold diluted by SP4 medium without antibiotics, plated onto SP4 agar plate containing 3 μg/mL puromycin and incubated at 37°C for four to five days. For the frozen competent cell protocol, cells were suspended with 140 μL of 0.1 M CaCl_2_, kept for 30 min on ice, and put into a −80°C deep freezer without additional glycerol or serum. The 70 μL of competent cell suspension in 1.5 mL tube was thawed on ice for 45 min.

### Colony PCR

To analyze transformants, colonies were picked up by 2 μL pipette tips through a digital microscope (Dino-Lite; AnMo Electronics, New Taipei, Taiwan), suspended 100 μL of SP4 medium containing 3 μg/mL puromycin and incubated at 37°C for one to two days. Cell suspensions were 10-fold diluted by ultra-pure water used as PCR templates. The PCR mixture contains 1 μL of template, 3 μM of primers (Puro-F1: 5’-ATG ACT GAA TAT AAA CCT ACT GTT AGA T-3’, and Puro-R1: 5’-TTA AGC ACC AGG TTT TCT AGT CAT ACA C-3’) and half volume of KOD One 2 × PCR master mix (TOYOBO, Osaka, Japan). The amplification cycle was performed by a thermal cycler (Dice Touch; TaKaRa, Shiga, Japan). Target bands were visualized by 2% agarose gel electrophoresis with GelRed staining and recorded by a gel imager (GelDoc; Bio-Rad, California, U.S.).

### Optical microscopy and Image analysis

Intact and competent cells were prepared from culture at pH 7.2 and 7.3. Intact cell culture and competent cells solutions were observed by phase-contrast optical microscopy using an inverted microscope (IX71; Olympus, Tokyo, Japan). Cell images were recorded by a complementary metal-oxide-semiconductor camera (DMK33UP5000; The Imaging Source, Bremen, Germany). Cell image intensity and diameter were measured by Analyze Particles macro in ImageJ 1.54f (https://imagej.net/ij/index.html).

## RESULTS

### Changes in the culture medium during growth of JCVI-syn3B

We collected basic data on the growth of JCVI-syn3B in SP4 medium, including pH, color, light absorbance, and viable cell count (CFU) at each growth stage. The growth of mycoplasma cells can be monitored by the color change of culture due to the presence of a pH indicator phenol red (16, 32). When JCVI-syn3B was grown in SP4 medium, the color of culture changed from red to orange or yellow (Supplementary Fig. S2A). To investigate the correlation between the color, pH, and absorbance at 560 nm of SP4 medium, we prepared SP4 media with various pH and measured absorbance at 560 nm. Gradual color changes from red to yellow with a decrease in pH were observed, and a strong correlation within the range of pH 7.6‒6.1 was found (R^2^ = 0.99) (Supplementary Fig S2B, C and Supplementary Table S2). To measure the CFU under various conditions, we prepared cells at different growth stages, measured their pH, and then diluted and plated them onto SP4 agar plates to count the number of colonies. The CFU increased from pH 7.4 to 7.0, remained stable from pH 7.0 to 6.5, and then gradually decreased from pH 6.5 (Supplementary Fig S2D and Supplementary Table S3).

In addition, we obtained a growth curve of JCVI-syn3B cells in SP4 medium using a plate reader by measuring absorbance at 560 nm continuously. A typical growth curve, including the lag phase, exponential growth phase, and stationary phase, was obtained. (Fig. 2). Since absorbance at 560 nm was correlated with pH within the range of pH 7.6‒ 6.1, we converted the absorbance to pH, and plotted CFUs on the growth curve (Fig. 2). As a result, CFU was observed to increase faster than changes in pH, and the observed growth stages could be divided into three phases: I, II, and III. Phase I was an initial phase where the number of cells increases, and absorbance and pH change rapidly. Phase II was a next phase where pH continued to become acidic while CFU reached a peak and entered the stationary state. In phase III, the culture became slightly more acidic, but the CFU decreased and the cells would enter the death phase (Fig. 2).

**FIGURE 2.**
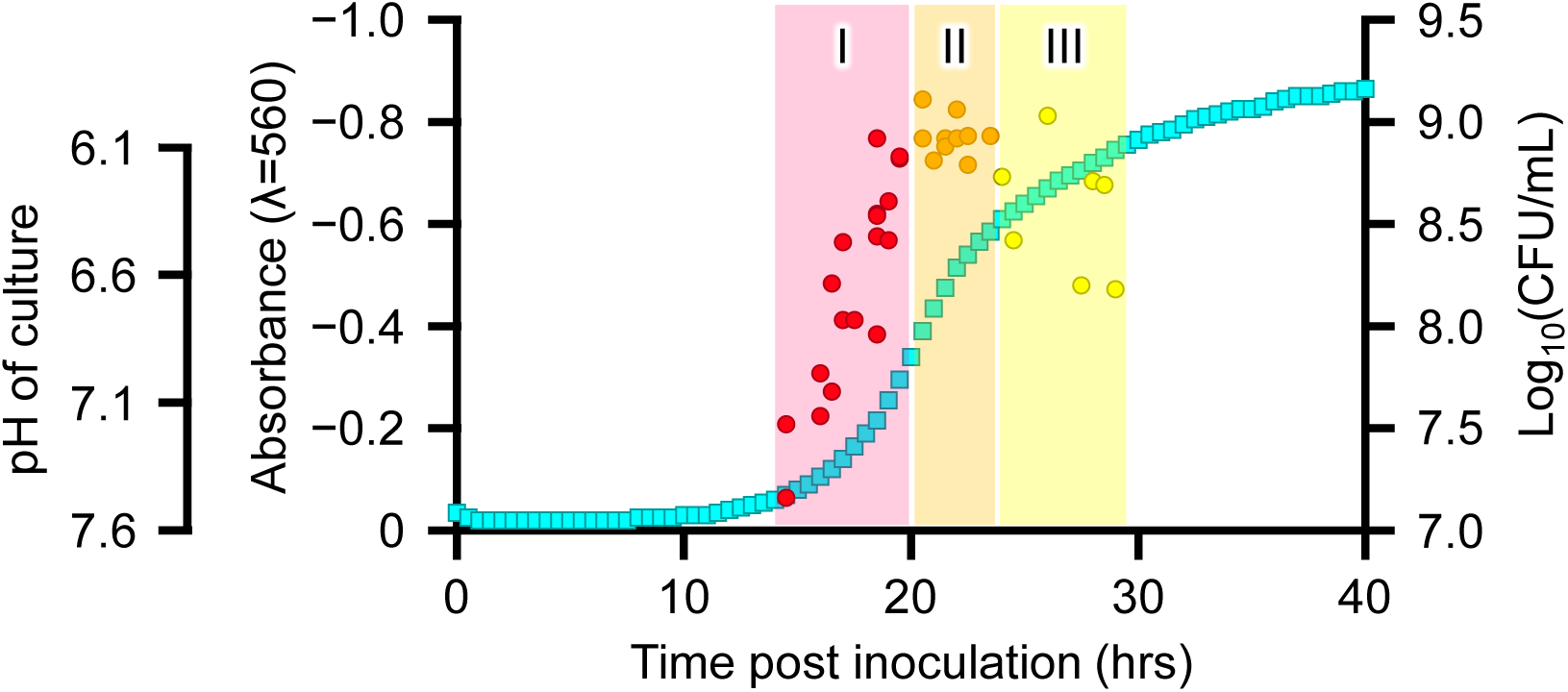
Growth phase of JCVI-syn3B. Growth of JCVI-syn3B in SP4 medium was monitored by light absorbance at 560 nm and is shown as light blue squares. The pH and absorbance at 560 nm in the SP4 media were correlated within the range of pH 7.6‒6.1, and pH is also shown on the left. The number of cells in cultures (colony forming units: CFU) was shown as circles. The axis for CFU is shown on the right. The growth phase after the lag phase could be divided into three phases: I, II, and III. In phase I, CFU increased rapidly, and pH also changed rapidly. In phase II, pH continued to become acidic while CFU reached a peak and entered the stationary state. In phase III, pH slightly continued to become acidic, but the CFU decreased and the cells entered the death phase.

### Optimization of transformation procedures

Based on previously reported PEG-mediated transformation methods for the minimal cells (16, 19, 26), we tried to refine procedures for development of a robust and highly efficient method. First, we investigated the transformation efficiency of recipient (competent) cells at different growth stages. JCVI-syn3B cells were grown and the growth stages were monitored by pH. Cultured cells were diluted and plated onto SP4 agar plate for CFU measurements, and also transformed with 100 ng of pSD128 and pSD131 plasmids by PEG-mediated transformation methods. Number of colonies on selection plates were counted to calculate transformation efficiency of individual cultures. The maximum transformation efficiency was obtained with higher pH: 5.4 × 10^-3^ and 3.1 × 10^-2^ (transformants/CFU/μg plasmid DNA) for pSD128 and pSD131 plasmids, respectively, at pH 7.31 (Fig. 3 and Supplementary Table S4). Transformation efficiency decreased when cells in the later phase were used: 7.0 × 10^-6^ and 1.1 × 10^-5^ for pSD128 and pSD131 plasmids, respectively, at pH 6.61. To confirm transformation, eight colonies from pSD128 and pSD131 transformants were picked and tested by colony PCR. All tested colonies showed the expected band in electrophoresis gels, suggesting all transformants were positive (Supplementary Fig. S3). These transformation efficiencies, especially when cells in the early phase were used, were extremely high compared to the previously reported transformation efficiencies of several *Mycoplasma* species (24, 33). To further modify the protocol, we tested methods to shorten the recovery time after PEG treatment. For the transformation of minimal cells described above, transformants were incubated for 3 hours in SP4 liquid medium before spreading on plates. To investigate the effect of recovery time for transformation efficiency, we compared transformation efficiency of transformants recovered for 1 and 2 hours. Transformation efficiencies using cells in exponential phase at pH 7.26 were 4.5 × 10^-3^ and 5.9 × 10^-3^ (transformants/CFU/μg plasmid DNA) with recovery times of 1 and 2 hours for pSD128, and 3.7 × 10^-2^ and 2.6 × 10^-2^ for 1 and 2 hours for pSD131, respectively (Supplementary Table S4). These efficiencies were not significantly different compared to efficiencies from experiments using with a 3-hour recovery time. Even transforming JCVI-syn3B cells that had grown longer and were no longer at a growth stage for optimal transformation 3-hour recovery times were no better than 1-hour recovery times.

**FIGURE 3.**
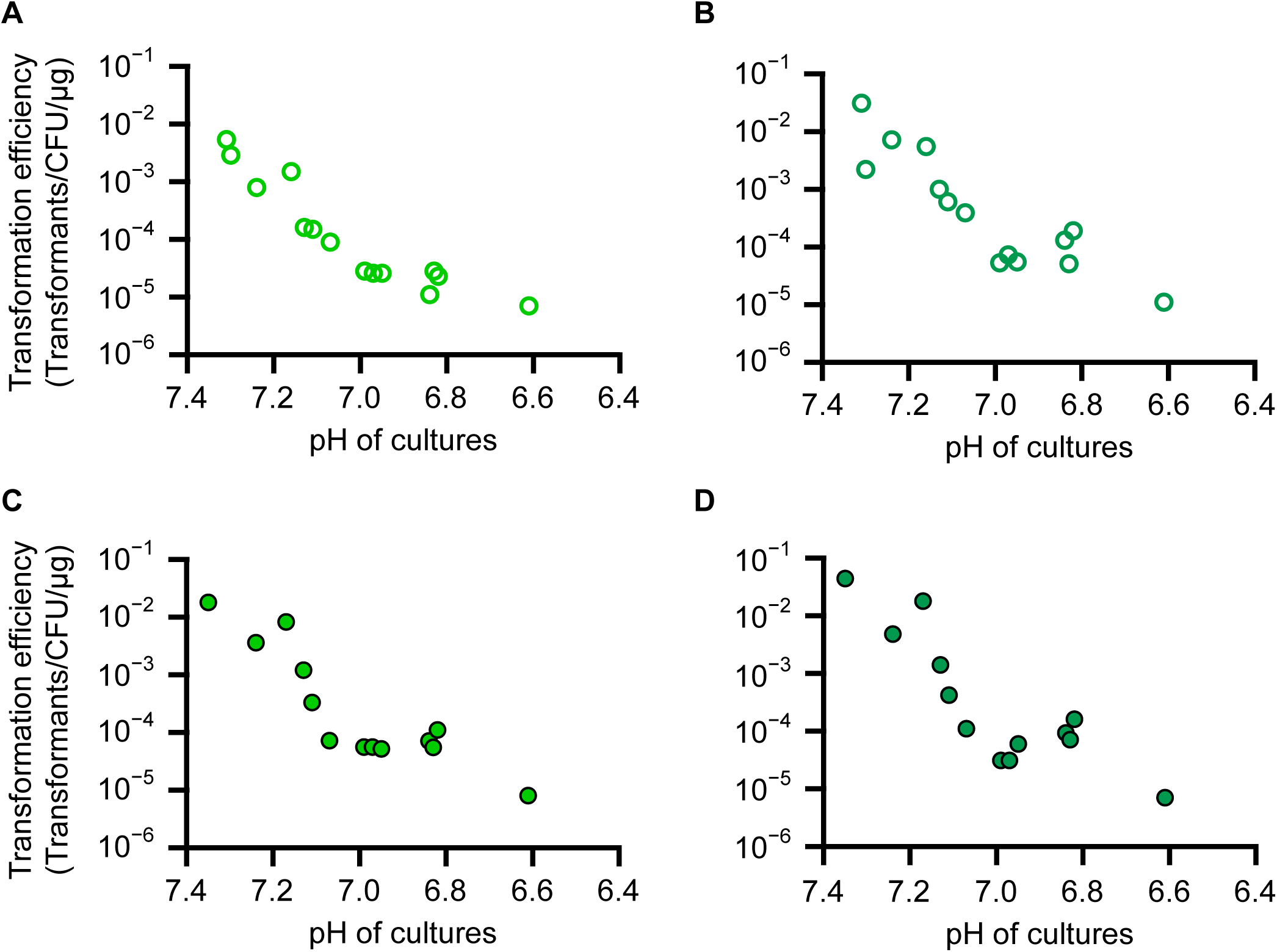
Transformation efficiency of JCVI-syn3B in various conditions. Transformation efficiencies using 100 ng of pSD128 (A) and pSD131 (B), and 10 ng of pSD128 (C) and pSD131 (D) are plotted as circles with pH of cultures. Transformation efficiencies of short recovery protocol using 100 ng plasmids are also plotted as open squares (2-hour recovery) and open triangles (1-hour recovery) in panels A and B.

Next, we tested transformations using a smaller amounts of plasmid DNA. Our initial experiments using 100 ng of plasmid DNA resulted in thousands to tens of thousands of colonies in the experiment described above (Supplementary Table S4). When the amount used was lowered to 10 ng, the transformation efficiency was similar to that obtained with 100 ng. We obtained approximately one-tenth the number of colonies, ranging from a few hundred to around 2000 colonies per experiment (Fig. 3C, D and Supplementary Table S4). The transformation efficiencies were 1.8 × 10^-2^ and 4.4 × 10^-2^ (transformants/CFU/μg plasmid DNA) for pSD128 and pSD131 plasmids at pH 7.35, and 8.0 × 10^-6^ and 7.0 × 10^-6^ at pH 6.61, respectively. Even using only 10 ng, a sufficient number of transformants could be obtained.

Thus using our optimized method, we obtained several hundred to several thousand transformants with less than 0.2 mL of cultured cells and 10 ng of plasmid, with shortened and simplified experimental procedures (Supplemental Table S4).

### Transformation using frozen competent cells

In *Escherichia coli* transformation, frozen competent cells are commonly used (34). However, to the best of our knowledge, there have been no reports using frozen competent cells for transformation of *Mycoplasma* species. To investigate the effects of freezing, we compared the number of living cells for intact culture, non-frozen competent cell solution, and frozen competent cell solution. Competent cell solutions were prepared and divided into two; one was plated for CFU measurement without freezing, and the other was frozen at −80°C. After one week or one month, the frozen solution was thawed on ice and plated for CFU measurement. The CFU decreased by 20% to 30% due to the competent cell preparation procedure and decreased by approximately one order of magnitude after freezing for 7 days (Supplementary Table S5).

To confirm the utility of frozen competent cells for transformation, we did transformation experiments using competent cells frozen for approximately one week (7‒ 12 days) and one month (30‒31 days). Even when freeze-thawed competent cells were used, a sufficient number of transformants were obtained in all the experiments tested, and the transformation efficiency was adequate (Fig. 4 and Supplementary Table S6). Transformation efficiency using one-month frozen competent cells was similar to that of one-week frozen competent cells, and both were lower by approximately one order of magnitude compared to non-frozen competent cells (Fig. 4, and Supplementary Table S4 and S6). The differences in transformation efficiency between frozen and non-frozen competent cells was likely attributable to cell death caused by freezing, as mentioned above (Supplementary Table S5). The relationship between the pH of the culture and transformation efficiency showed a similar tendency for both frozen and non-frozen cells: the transformation efficiency of frozen competent cells also decreased with the pH of the culture, as observed in the non-frozen competent cells (Fig. 3 and 4).

**FIGURE 4.**
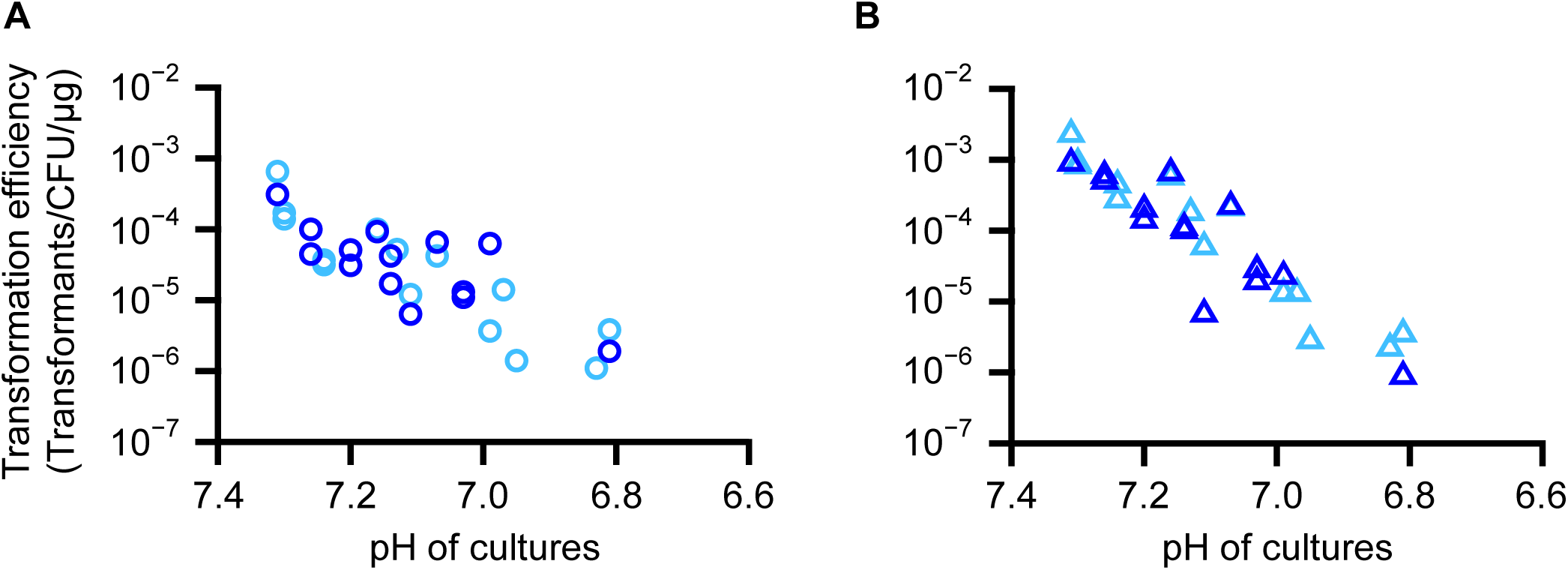
Transformation efficiency using frozen competent cells. Transformation efficiency using one-week or one-month frozen cells are plotted as light blue and blue, respectively. For all experiments, 100 ng plasmids were used. (A) pSD128, (B) pSD131.

To examine the state of cells during the preparation procedure of frozen competent cells, we conducted microscopic observations. Intact cells showed various cell morphologies. Most of cells showed round shape but the others were connected like chain or spread tentacles. Competent cells with CaCl_2_ showed round shape. (Supplementary Fig S4A, B). The freeze-thawed competent cells exhibited substantial variations in cell length, either becoming smaller or larger, indicating a more variability in cell dimension. Additionally, the image density of many cells was significantly reduced, with *p* = 6.5 × 10^-6^ (Supplementary Fig S4C, D). This might reflect cell damage caused by the freeze-thaw process in the competent state and was consistent with the lower transformation efficiency observed with frozen competent cells.

## DISCUSSION

### Summary of this study

Here, we optimized the PEG-mediated transformation method for the minimal cell JCVI-syn3B and successfully developed a robust and highly efficient transformation method. The newly established method using frozen competent cells will likely be useful for others working with JCVI-syn3B. We also obtained basic data on the pH, color of medium, light absorbance, and number of viable cells (CFU) that are used in PEG mediated transformation. During this process, we found that the JCVI-syn3B exponential growth phase can be divided into two distinct phases. Higher transformation efficiency was observed at the early-exponential phase. These methods and data would be useful for further utilization of the minimal cells.

### Growth stage of JCVI-syn3B

We examined the pH, color and absorbance of media, CFU, and transformation efficiency during cultivation. Through these processes, several findings were obtained as follows.

First, the pH of SP4 medium with phenol red could be monitored by light absorbance at 560 nm. Since absorbance at 560 nm correlated with pH within the pH range of 7.6‒ 6.1, absorbance at 560 nm alone can serve as a pH indicator of SP4 medium. In mycoplasmas, absorbance at 430 / 560 nm was usually used to monitor cell growth (16, 32), and absorbance at 430 nm might emphasize the growth curve. Highly efficient transformation was observed at the early-exponential phase in this study, so monitoring the cell state at 560 nm would be sufficient for efficient transformation purpose.

Second, CFU increased faster than changes in pH (Fig. 2). The growth stages observed by pH or absorbance could be divided into three phases: I, II, and III. Phase I was an initial phase where the number of cells increases, and absorbance and pH change rapidly. Phase II was a next phase where pH continued to become acidic while CFU reached a peak and entered the stationary state. In phase III, the media slightly continued to become acidic, but the CFU decreased and the cells entered the death phase (Fig. 2) The pH of culture would depend on intracellular metabolism: JCVI-syn3.0 could metabolize sugars like glucose to make ATP through the glycolysis pathway, and secrete mainly acetate outside the cells (21). Thus, the pH of the culture continues to decrease during growth, becoming more acidic, sometimes not correlated with cell division. The observation that pH did not always reflect the number of cells or cell stage would be logical. Particularly after the phase II, the pH continued to become acidic even as the number of cells remained the same or decreased because intracellular metabolism might have continued after cell division stopped.

Third, the transformation efficiency was the highest in the early-exponential stage, the early-phase I. Notably, in this phase, although the number of recipient cells was lower since cells were not fully grown, using cells in this phase resulted in a high number of transformants (Supplementary Fig. S4). This situation would be similar to the preparation of chemically competent *E. coli* cells: although the exponential phase typically continues until OD_600_ reaches around 0.7‒1.0 in *E. coli*, the transformation efficiency is higher when competent cells are prepared at around OD_600_ = 0.2 (34), and many protocols for the preparation of *E. coli* competent cells follow this method. The higher transformation efficiency using cells that have just entered the exponential growth phase might be a common feature in both *E. coli* and *Mycoplasma*.

These data would be useful for utilizing JCVI-syn3B cells; however, extension of these findings to other strains needs should be considered with caution. For example, the optimal pH might vary among *Mycoplasma* species. For the effective genome transplantation into *Mycoplasma capricolum*, closely related to the minimal cells, cultivation of the recipient cells up to pH 6.2 has been reported (9).

In the case of *E. coli* growth in LB medium, sugars are exhausted during the early-stationary phase (35), leading to the release of acetate and acetyl-phosphate outside the cell and a temporary decrease in the pH of the culture. Subsequently, amino acids are used as carbon sources, and then secreted acetate is taken up from the culture to produce ATP through the TCA cycle, resulting in an increase in pH (36, 37). Since mycoplasmas lack the TCA cycle, a similar metabolic network as *E. coli* was not present, making it difficult to compare the relationship between intracellular conditions and the pH of the medium.

### High efficiency of the method developed in this study

In various *Mycoplasma* and other Mollicutes species, transformation procedures have been reported to elucidate their gene functions or introducing new genes. Usually, PEG-mediated methods or electroporation are used for *Mycoplasma* transformation. Most of which required large amounts of plasmids and freshly prepared recipient cells. For example, in the PEG-mediated methods, 3 μg of plasmid was required for *Mycoplasma bovis* (27), and 10 μg for *Mycoplasma pulmonis* and *Mycoplasma hominis* (25, 33). In the electroporation methods, 10 μg was required for *Mycoplasma mobile* (38), 30 μg for *Mycoplasma pneumoniae* (39), and 10 µg for *Mycoplasma capricolum, M. mycoides, Spiroplasma citri*, and *M. pulmonis* (14, 40). Here, we developed a method that yields hundreds of transformants using only 10 ng of plasmid DNA and less than 0.2 mL of cultured recipient cells (Fig. 3C, D and Supplementary Table S4). Even when recipient cells in the stationary phase were used, around ten to hundred colonies were obtained. The minimal cell was created based on *M. mycoides* and is phylogenetically distant from clades such as *M. hominis* and *M. pneumoniae*, making direct comparison of transformation efficiencies with these *Mycoplasma* species difficult. Additionally, in the genome transplantation using *M. capricolum* as the recipient cell, the transplantation efficiency was reported to be affected by genome methylation and the activity of restriction enzymes (9, 41). The minimal cell was designed to lack genes for restriction enzymes and nucleases during the genome streamlining process (10), so transformation using the minimal cell as a recipient is expected to be highly efficient. Thus, the method developed in this study would be a robust, simple, and highly efficient transformation method for the minimal cells.

A slight difference was observed between the landing pad and *oriC* plasmids. When using 100 ng, *oriC* plasmids were more efficient than landing pad plasmids, but there was no difference when using 10 ng (Supplementary Table S4). The reason for this was unclear, but it might be influenced by impurities in the plasmid extract or the supercoiled state of the plasmids.

Previously reported transformation efficiencies of mycoplasmas were generally not high. For example, the transformation efficiency of the PEG-mediated method for *M. hominis* and *Mycoplasma arthritidis* were reported to be 2.3 × 10^-9^ and 1.2 × 10^-7^ (transformants/CFU/μg plasmid DNA), respectively (24, 33). In *M. mycoides* and *M capricolum*, the highest transformation efficiency with *oriC* plasmids was reported to be 6.0 × 10^−5^ (transformants/CFU/μg plasmid DNA) (14). These methods required large amounts of plasmid DNA and recipient cells. The maximum transformation efficiency obtained for JCVI-syn3B in this study was 4.4 × 10^-2^ (transformants/CFU/μg plasmid DNA) (Supplementary Table S4), allowing for sufficient colony numbers even with tiny amounts of plasmid DNA and recipient cells. The newly developed method was expected to be useful for future experiments requiring high transformation efficiency, such as library construction and random mutagenesis.

### Frozen competent cell of JCVI-syn3B

Preservation of JCVI-syn3B cells in a competent state at −80°C was also developed in this study (Supplementary Fig S4 and Supplementary Table S5). This eliminates the need to prepare new competent cells for each transformation experiment and allows for the storage of optimal competent cells in large quantities. Although long-term storage was not tested, the cells remained stable and maintained their competent state for at least one month (Fig. 4 and Supplementary Table S6). The transformation efficiency of the frozen competent cells was reduced by a factor of 10 compared to freshly prepared cells (Fig. 3 and 4), but the number of colonies obtained per experiment ranged from several dozen to several hundred, which would be sufficient for most experimental purposes. This method can make transformation of JCVI-syn3B and JCVI-syn3.0 minimal cells as fast and convenient as using frozen *E. coli* competent cells.

## Conclusion

The data and methods obtained in this study offer an improved more robust approach for installation of new genes into the minimized *M. mycoides* cells that are in wide use as model systems for a growing number of laboratories. It also suggests approaches for improving plasmid transformation of other mycoplasma species.

## DATA AVAILABILITY STATEMENT

The nucleotide sequence data for pSD128 and pSD131 were deposited in the DNA Data Bank of Japan (DDBJ) under the accession numbers LC823835 and LC823836.

## AUTHOR CONTRIBUTIONS

SK and MM conceived the study. SK, YO, JG and MM designed experimental plans and performed vector constructions. MM performed growth monitoring, transformation experiments and microscopic observation. SK and MM performed data analysis. SK, MM and TF wrote the paper. All authors edited the paper and approved the final version of the manuscript.

## FIGURE CAPTIONS

**SUPPLEMENTARY FIGURE S1.**

**Overview of the transformation procedures developed in this study.**

**SUPPLEMENTARY FIGURE S2.**

**pH and color change of SP4 media and number of JCVI-syn3B cells during cultivation.**

(A) Color changes during cultivation of JCVI-syn3B in a 1.5 mL tube. The left and right tubes are before and after cultivation. The pH of each culture is also shown. (B) Color change of SP4 medium under various pH conditions. The series of SP4 media were prepared by addition of HCl. The pH of each medium is shown at upper of each well. (C) Light absorbance at 560 nm in a series of SP4 media at various pH. The pH and absorbance were linearly correlated within the range of pH 7.6‒6.1 (R^2^ = 0.99). (D) The number of viable cells (CFU) at various pH of cultures.

**SUPPLEMENTARY FIGURE S3.**

**Colony PCRs to confirm transformation of JCVI-syn3B cells.**

The puromycin resistance gene was amplified. Lane 1: positive control (JCVI-syn3B strain containing the puromycin resistance gene). Lane 2: negative control (wild type JCVI-syn3B strain). Lanes 3‒10 and 11‒18 were transformants with pSD128 and pSD131, respectively.

**SUPPLEMENTARY FIGURE S4.**

**Morphology of JCVI-syn3B competent cells.**

Phase-contrast micrographs of (A) intact JCVI-syn3B cells, (B) non-frozen competent JCVI-syn3B cells, and (C) frozen competent JCVI-syn3B cells that were kept at −80°C for 7 days and then thawed. (D) Diameter and image density of individual cells were measured and plotted. A total of 105 and 111 cells from non-frozen and frozen competent cells were measured and plotted in red and blue, respectively. The legend for image density is shown on the right. Dashed gray lines indicate the maximum and minimum diameter of non-frozen competent cells. The mean diameter of the cells, mean image density, and their significance test (*t*-test) are shown in Supplementary Table S6.

**SUPPLEMENTARY FIGURE S5.**

**Summary of growth rate, number of cells (CFU), and transformation efficiency of JCVI-syn3B.**

Three kinds of data are overlaid: the growth rate (absorbance at 560 nm, squares in light blue) and CFU (filled circles) of JCVI-syn3B cells are the same as in Figure 2, and the transformation efficiencies are the same as in Figure 3. Only the data around the exponential phase are displayed. The transformation efficiencies with 100 ng of pSD128 and pSD131 plasmids are shown in open circles in light green and dark green, respectively. There are three different *X*-axis scales, each labeled with colors similar to the data plots. High transformation efficiency was observed in the early-exponential phase.

**SUPPLEMENTARY TABLE S1**

Recipe of SP4 medium.

**SUPPLEMENTARY TABLE S2.**

pH and light absorbance at 560 nm of SP4 medium.

**SUPPLEMENTARY TABLE S3.**

Number of cells (CFU) of JCVI-syn3B in cultured SP4 media.

**SUPPLEMENTARY TABLE S4.**

Transformation efficiency of JCVI-syn3B.

**SUPPLEMENTARY TABLE S5.**

Changes of CFU during freezing procedures.

**SUPPLEMENTARY TABLE S6.**

Transformation efficiency of frozen JCVI-syn3B competent cell.

**SUPPLEMENTARY TABLE S7.**

Mean diameter and image density of JCVI-syn3B competent cells by microscopy.

**SUPPLEMENTARY TABLE S8.**

Individual data for the diameter and image density of JCVI-syn3B competent cells by microscopy.

