## Supplementary Figure for "Robust and highly efficient transformation method for a minimal Mycoplasma cell"

#### Overview of this method

Less than 0.2 mL cultured cell + 10 ng plasmid DNA

↓

800-6,000 colonies

#### Procedures

1.0 mL cultured cell

↓

Collect cell, wash by 1.0 mL of S/T buffer

↓

Centrifuge, discard supernatant

↓

Suspend the cells in 140 μL of 0.1 M $CaCl_2$, incubate for 30 min on ice

↓

(optional) Cells can be stocked at −80°C

↓

Put 3 μL of plasmid DNA and 17 μL of the cell solution in a new tube, mix

↓

Incubate for 10–15 min

↓

Add 133 μL of 70% PEG6000, mix, incubate for 2 min

↓

Add 800 μL of S/T buffer, disperse the cells

↓

Centrifuge, discard supernatant

↓

Add 500 μL of medium without antibiotics, incubate for 1–3 hours at 37°C

↓

Plate on selective agar medium, incubate at 37°C

### Supplementary Figure S1

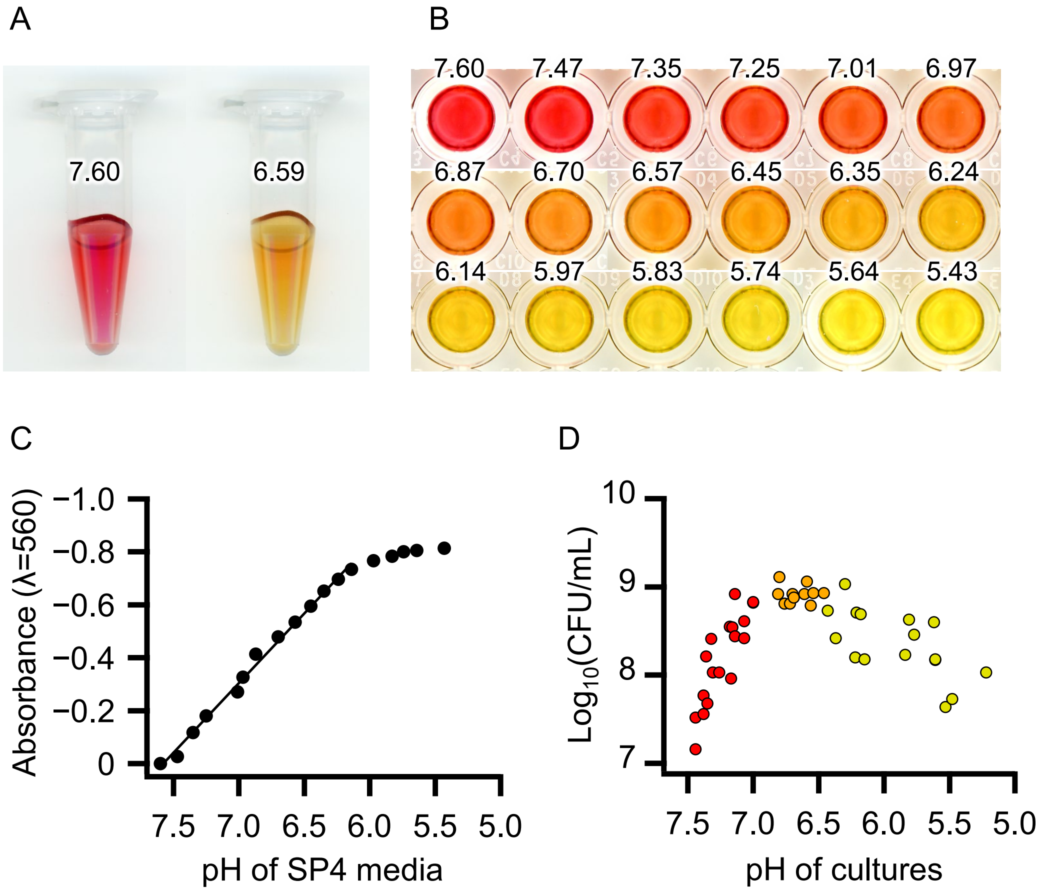


### Supplementary Figure S2

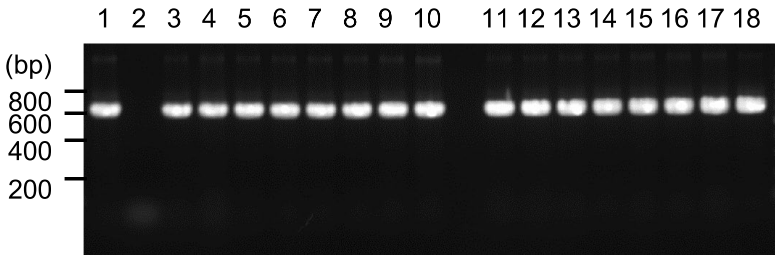


### Supplementary Figure S3

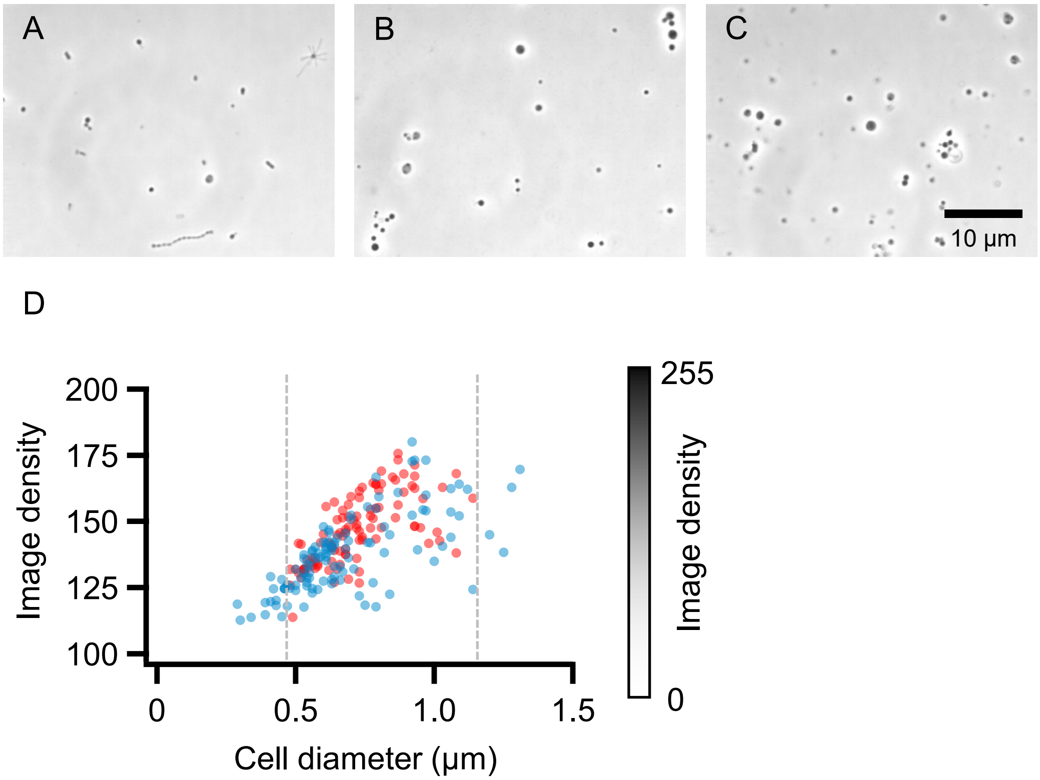


### Supplementary Figure S4

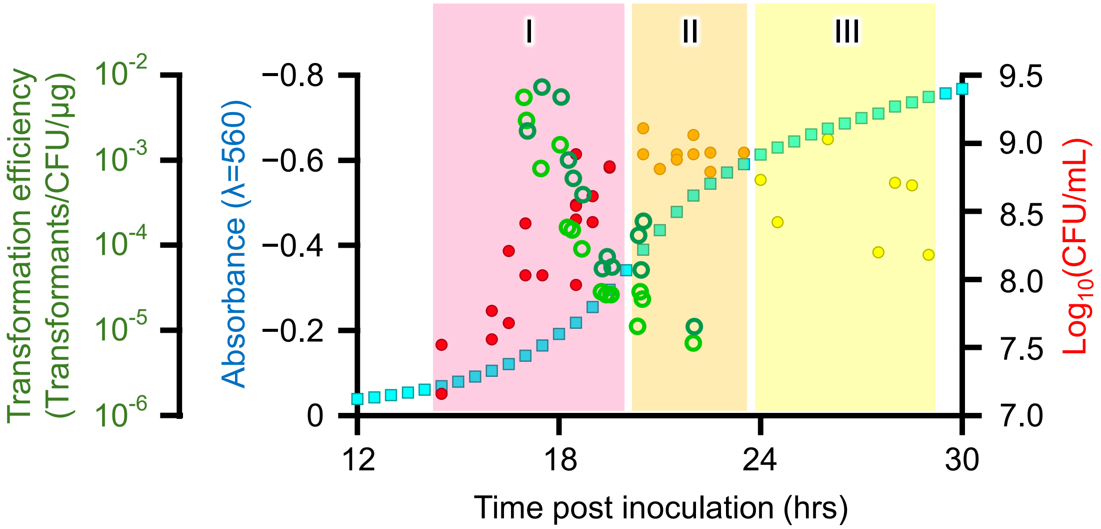


### Supplementary Figure S5
