## Supplementary Table for "Robust and highly efficient transformation method for a minimal Mycoplasma cell"

### Supplementary Table S1.
### Recipe of SP4 medium

| Basal Medium | Amounts for 1 L |
|---|---|
| Mycoplasma Broth Base | 3.5 g |
| Bacto Tryptone | 10 g |
| Bacto Peptone | 5.3 g |
| Agar (only for plate) | 10 g |
| Distilled water | 600 mL |
| 4 M KOH | 700 µL |
| | Sterilize at 121°C for 15 minutes. |
| **Sterile supplements** | **Amounts for 1 L** |
| 20% D-Glucose | 25 mL |
| CMRL 1066 (10×) | 50 mL |
| 7.5% Sodium Bicarbonate | 14.6 mL |
| 200 mM L-Glutamine | 5 mL |
| 25% Fresh Yeast extract Solution | 35 mL |
| 2% Yeastolate | 100 mL |
| Fetal Bovine Serum (Heat inactivated) | 170 mL |
| Penicillin G (400,000 units/mL) | 2.5 mL |
| 1% Phenol red | 1.5 mL |

### Supplementary Table S2.
### pH and $OD_{560}$ absorbance of SP4 medium

| pH* | $OD_{560}$** |
|---|---|
| 7.6 | 0 |
| 7.47 | -0.03 |
| 7.35 | -0.12 |
| 7.25 | -0.18 |
| 7.01 | -0.27 |
| 6.97 | -0.33 |
| 6.87 | -0.41 |
| 6.7 | -0.48 |
| 6.57 | -0.53 |
| 6.45 | -0.60 |
| 6.35 | -0.65 |
| 6.24 | -0.70 |
| 6.14 | -0.73 |
| 5.97 | -0.77 |
| 5.83 | -0.78 |
| 5.74 | -0.80 |
| 5.64 | -0.80 |
| 5.43 | -0.81 |

*pH of fresh SP4 medium for cultivation is 7.6.

**$OD_{560}$ absorbance of fresh SP4 was sabtracted.

### Supplementary Table S3.
### Number of cells (CFU) of JCVI-syn3B in cultured SP4 media

| pH of cultures | No. cells (CFU)/mL | $Log_{10}$(CFU/mL) |
|---|---|---|
| 7.44 | 3.3.E+07 | 7.52 |
| 7.44 | 1.4.E+07 | 7.16 |
| 7.38 | 5.8.E+07 | 7.77 |
| 7.38 | 3.6.E+07 | 7.56 |
| 7.36 | 1.6.E+08 | 8.21 |
| 7.35 | 4.8.E+07 | 7.68 |
| 7.32 | 2.6.E+08 | 8.41 |
| 7.31 | 1.1.E+08 | 8.03 |
| 7.26 | 1.1.E+08 | 8.03 |
| 7.18 | 3.6.E+08 | 8.55 |
| 7.17 | 9.2.E+07 | 7.96 |
| 7.16 | 3.5.E+08 | 8.54 |
| 7.14 | 8.3.E+08 | 8.92 |
| 7.14 | 2.8.E+08 | 8.44 |
| 7.07 | 2.6.E+08 | 8.42 |
| 7.07 | 4.1.E+08 | 8.61 |
| 7.00 | 6.7.E+08 | 8.82 |
| 7.00 | 6.8.E+08 | 8.83 |
| 6.81 | 8.3.E+08 | 8.92 |
| 6.80 | 1.3.E+09 | 9.11 |
| 6.76 | 6.4.E+08 | 8.81 |
| 6.72 | 6.5.E+08 | 8.81 |
| 6.70 | 8.3.E+08 | 8.92 |
| 6.69 | 7.6.E+08 | 8.88 |
| 6.61 | 8.3.E+08 | 8.92 |
| 6.59 | 1.1.E+09 | 9.06 |
| 6.56 | 6.1.E+08 | 8.79 |
| 6.54 | 8.6.E+08 | 8.93 |
| 6.46 | 8.5.E+08 | 8.93 |
| 6.43 | 5.4.E+08 | 8.73 |
| 6.37 | 2.7.E+08 | 8.42 |
| 6.30 | 1.1.E+09 | 9.03 |
| 6.22 | 1.6.E+08 | 8.20 |
| 6.21 | 5.2.E+08 | 8.71 |
| 6.18 | 4.9.E+08 | 8.69 |
| 6.15 | 1.5.E+08 | 8.18 |
| 5.84 | 1.7.E+08 | 8.23 |
| 5.81 | 4.3.E+08 | 8.63 |
| 5.77 | 2.9.E+08 | 8.46 |
| 5.62 | 4.0.E+08 | 8.60 |
| 5.61 | 1.5.E+08 | 8.17 |
| 5.61 | 1.5.E+08 | 8.18 |
| 5.53 | 4.4.E+07 | 7.64 |
| 5.48 | 5.4.E+07 | 7.73 |
| 5.22 | 1.1.E+08 | 8.03 |

### Supplementary Table S4.
### Transformation efficiency of JCVI-syn3B

| pH of cultures | No. cells (CFU)/mL | pSD128 (ng) | pSD131 (ng) | Recovery (hrs) | No. Transformants with pSD128* | No. Transformants with pSD131* | pSD128 Efficiency (Transformants/CFU/µg) | pSD131 Efficiency (Transformants/CFU/µg) |
|---|---|---|---|---|---|---|---|---|
| **100 ng plasmid** | | | | | | | | |
| 7.31 | 1.1.E+08 | 100 | 100 | 3 | 7010 | 40440 | 5.4.E-03 | 3.1.E-02 |
| 7.30 | 2.4.E+08 | 100 | 100 | 3 | 8600 | 6400 | 2.9.E-03 | 2.2.E-03 |
| 7.24 | 3.3.E+08 | 100 | 100 | 3 | 3200 | 29000 | 7.9.E-04 | 7.2.E-03 |
| 7.16 | 3.5.E+08 | 100 | 100 | 3 | 6150 | 23030 | 1.5.E-03 | 5.5.E-03 |
| 7.13 | 5.9.E+08 | 100 | 100 | 3 | 1130 | 7150 | 1.6.E-04 | 1.0.E-03 |
| 7.11 | 3.2.E+08 | 100 | 100 | 3 | 570 | 2360 | 1.5.E-04 | 6.1.E-04 |
| 7.07 | 2.6.E+08 | 100 | 100 | 3 | 290 | 1250 | 9.0.E-05 | 3.9.E-04 |
| 6.99 | 2.7.E+08 | 100 | 100 | 3 | 90 | 170 | 2.8.E-05 | 5.3.E-05 |
| 6.97 | 5.8.E+08 | 100 | 100 | 3 | 184 | 520 | 2.6.E-05 | 7.3.E-05 |
| 6.95 | 7.1.E+08 | 100 | 100 | 3 | 227 | 479 | 2.6.E-05 | 5.5.E-05 |
| 6.83 | 8.4.E+08 | 100 | 100 | 3 | 284 | 519 | 2.8.E-05 | 5.1.E-05 |
| 6.82 | 6.6.E+08 | 100 | 100 | 3 | 188 | 1520 | 2.3.E-05 | 1.9.E-04 |
| 6.61 | 8.3.E+08 | 100 | 100 | 3 | 70 | 110 | 7.0.E-06 | 1.1.E-05 |
| Average, phase 1 (pH 7.13-7.31) | | | | | 5218† | 21204† | 2.2.E-03 | 9.4.E-03 |
| Average, phase 2 (pH 6.61-7.11) | | | | | 238 | 866 | 4.7.E-05 | 1.8.E-04 |
| **Test of recovery time** | | | | | | | | |
| 7.26 | 1.1.E+08 | 100 | 100 | 2 | 7700 | 33700 | 5.9.E-03 | 2.6.E-02 |
| 7.15 | 2.9.E+08 | 100 | 100 | 2 | 410 | 2000 | 1.2.E-04 | 5.6.E-04 |
| 7.07 | 4.1.E+08 | 100 | 100 | 2 | 1944 | 3560 | 3.9.E-04 | 7.2.E-04 |
| 7.01 | 5.5.E+08 | 100 | 100 | 2 | 206 | 585 | 3.1.E-05 | 8.8.E-05 |
| 6.81 | 8.3.E+08 | 100 | 100 | 2 | 68 | 140 | 6.7.E-06 | 1.4.E-05 |
| 6.75 | 6.2.E+08 | 100 | 100 | 2 | 126 | 314 | 1.7.E-05 | 4.2.E-05 |
| 6.72 | 6.5.E+08 | 100 | 100 | 2 | 124 | 232 | 1.6.E-05 | 2.9.E-05 |
| 7.26 | 1.1.E+08 | 100 | 100 | 1 | 5900 | 48800 | 4.5.E-03 | 3.7.E-02 |
| 7.15 | 2.9.E+08 | 100 | 100 | 1 | 130 | 1000 | 3.7.E-05 | 2.8.E-04 |
| 7.07 | 4.1.E+08 | 100 | 100 | 1 | 1908 | 3328 | 3.9.E-04 | 6.8.E-04 |
| 7.01 | 5.5.E+08 | 100 | 100 | 1 | 149 | 404 | 2.2.E-05 | 6.0.E-05 |
| 6.81 | 8.3.E+08 | 100 | 100 | 1 | 8 | 8 | 7.9.E-07 | 7.9.E-07 |
| 6.75 | 6.2.E+08 | 100 | 100 | 1 | 26 | 267 | 3.5.E-06 | 3.6.E-05 |
| 6.72 | 6.5.E+08 | 100 | 100 | 1 | 68 | 112 | 8.6.E-06 | 1.4.E-05 |
| **10 ng plasmid** | | | | | | | | |
| 7.35 | 4.8.E+07 | 10 | 10 | 3 | 1060 | 2580 | 1.8.E-02 | 4.4.E-02 |
| 7.30 | 2.4.E+08 | 10 | 10 | 3 | 5660 | 6590 | 1.9.E-02 | 2.2.E-02 |
| 7.24 | 3.3.E+08 | 10 | 10 | 3 | 1470 | 1930 | 3.6.E-03 | 4.8.E-03 |
| 7.17 | 9.2.E+07 | 10 | 10 | 3 | 920 | 2010 | 8.2.E-03 | 1.8.E-02 |
| 7.13 | 5.9.E+08 | 10 | 10 | 3 | 850 | 1023 | 1.2.E-03 | 1.4.E-03 |
| 7.11 | 3.2.E+08 | 10 | 10 | 3 | 127 | 164 | 3.3.E-04 | 4.2.E-04 |
| 7.07 | 2.6.E+08 | 10 | 10 | 3 | 23 | 35 | 7.2.E-05 | 1.1.E-04 |
| 6.99 | 2.7.E+08 | 10 | 10 | 3 | 18 | 10 | 5.6.E-05 | 3.1.E-05 |
| 6.97 | 5.8.E+08 | 10 | 10 | 3 | 40 | 22 | 5.6.E-05 | 3.1.E-05 |
| 6.95 | 7.1.E+08 | 10 | 10 | 3 | 45 | 52 | 5.2.E-05 | 6.0.E-05 |
| 6.83 | 8.4.E+08 | 10 | 10 | 3 | 56 | 72 | 5.5.E-05 | 7.1.E-05 |
| 6.82 | 6.6.E+08 | 10 | 10 | 3 | 90 | 130 | 1.1.E-04 | 1.6.E-04 |
| 6.61 | 8.3.E+08 | 10 | 10 | 3 | 8 | 7 | 8.0.E-06 | 7.0.E-06 |
| Average, phase 1 (pH 7.13-7.31) | | | | | 1992† | 2826† | 1.0.E-02 | 1.8.E-02 |
| Average, phase 2 (pH 6.61-7.11) | | | | | 51 | 62 | 9.2.E-05 | 1.1.E-04 |

*These transformants (colonies) were obtained using 0.12 mL (17 µL amoung 140 µL of $CaCl_2$-treated cells) of cultured cells.

† These were significantly different from the average numbers of phase 2.

### Supplementary Table S5.
### Changes of CFU during freezing procedures.

| pH of cultures | Status | CFU/mL |
|---|---|---|
| 7.3 | Intact culture | 9.0.E+07 |
| | Non-frozen competent | 7.1.E+07 |
| | Frozen competent | 8.2.E+06 |
| 7.2 | Intact culture | 2.8.E+08 |
| | Non-frozen competent | 1.9.E+08 |
| | Frozen competent | 5.6.E+07 |

### Supplementary Table S6.
### Transformation efficiency of frozen JCVI-syn3B competent cell

| pH of cultures | No. cells (CFU)/mL | pSD128 (ng) | pSD131 (ng) | Recovery (hrs) | Days frozen | No. Transformants with pSD128* | No. Transformants with pSD131* | pSD128 Efficiency (Transformants/CFU/μg) | pSD131 Efficiency (Transformants/CFU/μg) |
|---|---|---|---|---|---|---|---|---|---|
| **1 week** | | | | | | | | | |
| 7.31 | 1.1.E+08 | 100 | 100 | 3 | 8 | 840 | 2990 | 6.5.E-04 | 2.3.E-03 |
| 7.30 | 2.4.E+08 | 100 | 100 | 3 | 7 | 410 | 2640 | 1.4.E-04 | 9.0.E-04 |
| 7.30 | 2.4.E+08 | 100 | 100 | 3 | 7 | 510 | 2410 | 1.7.E-04 | 8.2.E-04 |
| 7.24 | 3.3.E+08 | 100 | 100 | 3 | 7 | 150 | 1770 | 3.7E-05 | 4.4E-04 |
| 7.24 | 3.3.E+08 | 100 | 100 | 3 | 7 | 130 | 1090 | 3.2E-05 | 2.7E-04 |
| 7.16 | 3.5.E+08 | 100 | 100 | 3 | 8 | 430 | 2410 | 1.0.E-04 | 5.7.E-04 |
| 7.13 | 5.9.E+08 | 100 | 100 | 3 | 8 | 371 | 1247 | 5.2.E-05 | 1.8.E-04 |
| 7.11 | 3.2.E+08 | 100 | 100 | 3 | 7 | 40 | 360 | 1.0.E-05 | 9.3.E-05 |
| 7.11 | 3.2.E+08 | 100 | 100 | 3 | 12 | 48 | 228 | 1.2.E-05 | 5.9.E-05 |
| 7.07 | 4.1.E+08 | 100 | 100 | 3 | 10 | 208 | 1032 | 4.2.E-05 | 2.1.E-04 |
| 6.99 | 2.7.E+08 | 100 | 100 | 3 | 12 | 12 | 42 | 3.7.E-06 | 1.3.E-05 |
| 6.97 | 5.8.E+08 | 100 | 100 | 3 | 8 | 99 | 91 | 1.4.E-05 | 1.3.E-05 |
| 6.95 | 7.1.E+08 | 100 | 100 | 3 | 7 | 12 | 24 | 1.4.E-06 | 2.8.E-06 |
| 6.83 | 8.4.E+08 | 100 | 100 | 3 | 8 | 11 | 22 | 1.1.E-06 | 2.2.E-06 |
| 6.81 | 8.3.E+08 | 100 | 100 | 3 | 10 | 38 | 35 | 3.8.E-06 | 3.5.E-06 |
| **1 month** | | | | | | | | | |
| 7.31 | 1.1.E+08 | 100 | 100 | 3 | 30 | 400 | 1160 | 3.1.E-04 | 9.0.E-04 |
| 7.26 | 2.2.E+08 | 100 | 100 | 3 | 31 | 120 | 1320 | 4.5.E-05 | 5.0.E-04 |
| 7.26 | 2.2.E+08 | 100 | 100 | 3 | 33 | 270 | 1600 | 1.0.E-04 | 6.0.E-04 |
| 7.20 | 2.4.E+08 | 100 | 100 | 3 | 32 | 150 | 580 | 5.1.E-05 | 2.0.E-04 |
| 7.20 | 2.4.E+08 | 100 | 100 | 3 | 32 | 90 | 400 | 3.1.E-05 | 1.4.E-04 |
| 7.16 | 3.5.E+08 | 100 | 100 | 3 | 30 | 390 | 2730 | 9.3.E-05 | 6.5.E-04 |
| 7.14 | 2.5.E+08 | 100 | 100 | 3 | 30 | 53 | 298 | 1.7.E-05 | 9.7.E-05 |
| 7.14 | 2.5.E+08 | 100 | 100 | 3 | 32 | 128 | 334 | 4.2.E-05 | 1.1.E-04 |
| 7.11 | 3.2.E+08 | 100 | 100 | 3 | 30 | 25 | 26 | 6.4.E-06 | 6.7.E-06 |
| 7.07 | 4.1.E+08 | 100 | 100 | 3 | 30 | 324 | 1066 | 6.6.E-05 | 2.2.E-04 |
| 7.03 | 3.3.E+08 | 100 | 100 | 3 | 32 | 52 | 112 | 1.3.E-05 | 2.8.E-05 |
| 7.03 | 3.3.E+08 | 100 | 100 | 3 | 32 | 44 | 77 | 1.1.E-05 | 1.9.E-05 |
| 6.81 | 8.3.E+08 | 100 | 100 | 3 | 30 | 19 | 9 | 1.9.E-06 | 8.9.E-07 |

*These transformants (colonies) were obtained using 0.12 mL (17 μL amoung 140 μL of $CaCl_2$-treated cells) of cultured cells.

### Supplementary Table S7.
### Mean diameter and image density of JCVI-syn3B competent cells by microscopy

| | No. cells count | Mean Diameter* (μm) | Image density* |
|---|---|---|---|
| Fresh competent cell | 105 | 0.69 ± 0.22 | 137.7 ±14.8 |
| Frozen competent cell | 111 | 0.72 ± 0.15 | 146.5 ± 12.9 |
| *t* -test | | 0.20 | 6.5E-06 |
| *SD values are also shown. | | | |

### Supplementary Table S8.
### Individual data for the diameter and image density of JCVI-syn3B competent cells by microscopy.

| Fresh competent cells | | |
|---|---|---|
| No. | Diameter (μm) | Image density |
| 1 | 0.61 | 155.6 |
| 2 | 0.67 | 154.2 |
| 3 | 0.72 | 152.0 |
| 4 | 0.93 | 148.2 |
| 5 | 0.65 | 144.9 |
| 6 | 0.54 | 132.2 |
| 7 | 0.48 | 131.9 |
| 8 | 0.63 | 140.1 |
| 9 | 0.74 | 143.0 |
| 10 | 0.98 | 141.7 |
| 11 | 0.95 | 147.6 |
| 12 | 0.50 | 131.9 |
| 13 | 0.56 | 136.3 |
| 14 | 0.77 | 154.5 |
| 15 | 0.53 | 131.8 |
| 16 | 0.70 | 159.4 |
| 17 | 0.86 | 151.4 |
| 18 | 0.87 | 156.2 |
| 19 | 0.63 | 134.8 |
| 20 | 0.64 | 157.3 |
| 21 | 0.58 | 133.1 |
| 22 | 0.79 | 155.0 |
| 23 | 0.73 | 131.0 |
| 24 | 0.69 | 147.9 |
| 25 | 1.14 | 158.8 |
| 26 | 0.64 | 139.1 |
| 27 | 0.57 | 134.8 |
| 28 | 0.52 | 128.7 |
| 29 | 0.53 | 131.9 |
| 30 | 0.49 | 113.8 |
| 31 | 0.57 | 132.3 |
| 32 | 0.64 | 127.0 |
| 33 | 0.52 | 141.3 |
| 34 | 0.72 | 148.8 |
| 35 | 0.51 | 141.7 |
| 36 | 0.77 | 152.1 |
| 37 | 0.55 | 134.2 |
| 38 | 0.67 | 144.5 |
| 39 | 0.74 | 144.1 |
| 40 | 0.80 | 155.2 |
| 41 | 0.57 | 136.3 |
| 42 | 0.69 | 151.8 |
| 43 | 0.74 | 162.9 |
| 44 | 0.59 | 141.9 |
| 45 | 0.81 | 164.2 |
| 46 | 0.53 | 130.5 |
| 47 | 0.54 | 135.6 |
| 48 | 0.66 | 145.7 |
| 49 | 0.79 | 143.4 |
| 50 | 0.72 | 147.4 |
| 51 | 0.69 | 128.2 |
| 52 | 0.93 | 162.5 |
| 53 | 0.51 | 130.6 |
| 54 | 0.53 | 131.4 |
| 55 | 0.68 | 137.6 |
| 56 | 0.72 | 136.0 |
| 57 | 0.48 | 125.9 |
| 58 | 0.63 | 140.3 |
| 59 | 0.53 | 132.1 |
| 60 | 0.73 | 142.9 |
| 61 | 0.96 | 158.7 |
| 62 | 1.08 | 168.1 |
| 63 | 0.78 | 164.6 |
| 64 | 0.68 | 145.8 |
| 65 | 0.87 | 173.3 |
| 66 | 0.78 | 151.1 |
| 67 | 0.68 | 137.5 |
| 68 | 0.56 | 133.2 |
| 69 | 0.62 | 131.6 |
| 70 | 0.68 | 139.6 |
| 71 | 0.73 | 126.7 |
| 72 | 0.79 | 164.0 |
| 73 | 1.08 | 138.1 |
| 74 | 0.80 | 161.8 |
| 75 | 0.58 | 133.9 |
| 76 | 0.65 | 132.0 |
| 77 | 1.02 | 142.7 |
| 78 | 0.93 | 171.4 |
| 79 | 0.89 | 167.9 |
| 80 | 0.81 | 147.7 |
| 81 | 0.73 | 146.3 |
| 82 | 0.85 | 166.7 |
| 83 | 0.77 | 147.5 |
| 84 | 1.01 | 146.0 |
| 85 | 0.92 | 163.5 |
| 86 | 0.68 | 146.9 |
| 87 | 0.69 | 156.3 |
| 88 | 0.73 | 161.4 |
| 89 | 0.60 | 145.2 |
| 90 | 0.62 | 141.9 |
| 91 | 0.71 | 150.5 |
| 92 | 0.87 | 175.7 |
| 93 | 0.81 | 169.1 |
| 94 | 0.89 | 161.1 |
| 95 | 0.79 | 164.1 |
| 96 | 0.93 | 148.1 |
| 97 | 0.93 | 167.1 |
| 98 | 1.03 | 162.9 |
| 99 | 0.65 | 150.7 |
| 100 | 0.86 | 165.6 |
| 101 | 0.63 | 141.1 |
| 102 | 0.66 | 138.8 |
| 103 | 0.73 | 159.0 |
| 104 | 0.67 | 151.4 |
| 105 | 0.66 | 133.7 |

| Frozen competent cells | | |
|---|---|---|
| Cell_ID | Diameter (μm) | Image density |
| 1 | 0.53 | 130.4 |
| 2 | 1.31 | 169.7 |
| 3 | 0.53 | 136.0 |
| 4 | 0.60 | 148.0 |
| 5 | 0.92 | 180.1 |
| 6 | 0.61 | 137.8 |
| 7 | 0.56 | 138.6 |
| 8 | 0.92 | 152.4 |
| 9 | 0.96 | 154.4 |
| 10 | 0.47 | 126.1 |
| 11 | 1.12 | 162.2 |
| 12 | 0.93 | 173.2 |
| 13 | 0.58 | 136.0 |
| 14 | 0.62 | 145.9 |
| 15 | 0.55 | 130.7 |
| 16 | 0.61 | 135.2 |
| 17 | 0.50 | 131.9 |
| 18 | 0.92 | 172.7 |
| 19 | 0.43 | 118.3 |
| 20 | 0.39 | 114.7 |
| 21 | 0.63 | 128.1 |
| 22 | 0.56 | 123.0 |
| 23 | 0.56 | 124.7 |
| 24 | 0.70 | 150.9 |
| 25 | 0.68 | 145.6 |
| 26 | 1.09 | 164.1 |
| 27 | 1.06 | 144.0 |
| 28 | 0.69 | 142.7 |
| 29 | 0.82 | 148.0 |
| 30 | 0.76 | 155.5 |
| 31 | 1.06 | 162.4 |
| 32 | 1.03 | 140.6 |
| 33 | 0.54 | 125.6 |
| 34 | 0.47 | 118.0 |
| 35 | 0.70 | 152.4 |
| 36 | 0.79 | 154.9 |
| 37 | 0.71 | 132.1 |
| 38 | 0.65 | 143.9 |
| 39 | 0.60 | 127.4 |
| 40 | 0.50 | 124.0 |
| 41 | 0.64 | 141.0 |
| 42 | 0.67 | 130.9 |
| 43 | 0.45 | 114.0 |
| 44 | 1.00 | 135.0 |
| 45 | 0.87 | 160.9 |
| 46 | 0.63 | 126.5 |
| 47 | 0.66 | 127.8 |
| 48 | 0.97 | 173.2 |
| 49 | 0.61 | 139.4 |
| 50 | 0.80 | 159.4 |
| 51 | 0.58 | 138.1 |
| 52 | 0.62 | 146.8 |
| 53 | 0.45 | 128.1 |
| 54 | 0.61 | 136.6 |
| 55 | 1.28 | 162.9 |
| 56 | 0.41 | 129.1 |
| 57 | 0.64 | 142.0 |
| 58 | 0.82 | 138.2 |
| 59 | 0.29 | 118.8 |
| 60 | 0.68 | 139.7 |
| 61 | 1.09 | 152.1 |
| 62 | 0.63 | 139.0 |
| 63 | 0.76 | 142.1 |
| 64 | 1.20 | 145.0 |
| 65 | 0.84 | 145.0 |
| 66 | 0.61 | 141.7 |
| 67 | 0.97 | 154.0 |
| 68 | 0.62 | 142.5 |
| 69 | 0.48 | 124.8 |
| 70 | 0.41 | 119.7 |
| 71 | 0.60 | 142.2 |
| 72 | 1.25 | 138.3 |
| 73 | 0.39 | 119.4 |
| 74 | 0.79 | 166.7 |
| 75 | 0.57 | 140.4 |
| 76 | 0.55 | 128.7 |
| 77 | 0.55 | 133.3 |
| 78 | 0.46 | 124.7 |
| 79 | 0.56 | 135.5 |
| 80 | 0.64 | 129.1 |
| 81 | 1.06 | 153.6 |
| 82 | 0.50 | 125.9 |
| 83 | 0.69 | 136.3 |
| 84 | 0.63 | 140.0 |
| 85 | 0.58 | 124.3 |
| 86 | 0.75 | 118.4 |
| 87 | 0.56 | 139.0 |
| 88 | 0.79 | 117.8 |
| 89 | 0.53 | 137.4 |
| 90 | 1.14 | 124.3 |
| 91 | 0.54 | 133.6 |
| 92 | 0.94 | 139.3 |
| 93 | 0.54 | 128.9 |
| 94 | 0.46 | 124.7 |
| 95 | 0.59 | 136.6 |
| 96 | 0.54 | 127.0 |
| 97 | 0.73 | 121.9 |
| 98 | 0.34 | 113.8 |
| 99 | 0.52 | 128.8 |
| 100 | 0.42 | 124.6 |
| 101 | 0.30 | 112.7 |
| 102 | 0.97 | 160.0 |
| 103 | 0.64 | 133.0 |
| 104 | 0.46 | 124.7 |
| 105 | 0.77 | 141.9 |
| 106 | 0.61 | 140.2 |
| 107 | 0.78 | 126.8 |
| 108 | 0.43 | 120.2 |
| 109 | 0.53 | 117.6 |
| 110 | 0.84 | 122.5 |
| 111 | 0.66 | 133.0 |
